# Assigning Secondary Structure in Proteins using AI

**DOI:** 10.1101/2021.02.02.429329

**Authors:** Jisna Vellara Antony, Prayagh Madhu, Jayaraj Pottekkattuvalappil Balakrishnan

**Affiliations:** Department of Computer Science and Engineering, National Institute of Technology Calicut, Kerala 673601, India; Computer Science and Engineering Dept., Rajiv Gandhi Institute of Technology, Kottayam, India

**Keywords:** Protein Structure Assignment, Deep Learning, Fragment Library Creation, Convolutional Neural Networks, Protein Fragments, Protein Secondary Structures, Multi-Class Classifier

## Abstract

Knowledge about protein structure assignment enriches the structural and functional understanding of proteins. Accurate and reliable structure assignment data is crucial for secondary structure prediction systems. Since the ’80s various methods based on hydrogen bond analysis and atomic coordinate geometry, followed by Machine Learning, have been employed in protein structure assignment. However, the assignment process becomes challenging when missing atoms are present in protein files. Our model develops a multi-class classifier program named DLFSA for assigning protein Secondary Structure Elements(SSE) using Convolutional Neural Networks(CNN). A fast and efficient GPU based parallel procedure extracts fragments from protein files. The model implemented in this work is trained with a subset of protein fragments and achieves 88.1% and 82.5% train and test accuracy, respectively. Our model uses only C*α* coordinates for secondary structure assignments. The model is successfully tested on a few full-length proteins also. Results from the fragment-based studies demonstrate the feasibility of applying deep learning solutions for structure assignment problems.

## 1 Introduction

Pauling and Corey identify the existence of regular substructures namely, *α helices*(H) and *β sheets*(E), in protein molecules[1]. Irregular curves connecting these regular structures are called coils(C)[2][3][4]. This three-state classification extends to a finer eight state classification that includes the states, viz. 310 helices (G), *α helices*(H), *π helices* (I), *β strands* (E), *β bridges* (B), turns (T), bends (S), and others (C)[5]. Among these eight states of secondary structure, some states occur rarely. Protein structure assignment is the process of associating secondary-structure information into experimentally determining coordinates of a protein. These secondary structure information has contributed to structural and computational chemistry, viz—protein structure modelling, protein design, structure comparisons, classifications, and visualizations. Protein structure prediction is a hard task, involving many degrees of freedom. Protein secondary structures are a simpler representation of its tertiary structure. Prior knowledge about the secondary structure reduces the complexity associated with the modelling and design tasks. Most protein structure modelling systems use this secondary structure information at their initial steps, as it cuts down the conformational search space substantially, and thereby accelerating the whole prediction process[6][7][8]. These secondary structure prediction systems require structure assignment data that serves as ground truth for training the models.

Structure assignment was initially done manually by experts in the area, using visual inspection techniques, which often create discrepancies that lead to the process’s automation. Later on, scientists implement various computational structure assignment tools. Most of these assignment programs operate on hydrogen bond and Cartesian coordinate analysis, and a few works by machine learning approaches. DSSP(Dictionary of Protein Secondary Structures)[9] and STRIDE(STructural IDEntification) are two gold standards in protein structure assignment. DSSP works on fine-grained protein structures and confirms a hydrogen bond when the electrostatic energy-E between each interacting pair is less than *−*0.5 kcal/mole.

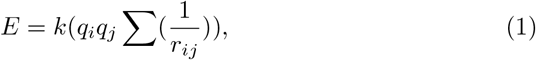

Where *q*_*i*_, *q*_*j*_ represents the charges on atoms separated by a distance *r*_*ij*_ and, k is the Coulomb constant. By analyzing the hydrogen bonding patterns obtained, DSSP annotates the sequence with the secondary structure information. As it is practically impossible to obtain fine-grained information for all protein structures in repositories, a fast and accurate method that performs well with C*α* positions is preferable. STRIDE[10] examines the phi-psi angle information along with hydrogen bond analysis data. PSSC[11], works on DSSP’s output and provides a more advanced characterization of secondary structures.

SECSTR[12] locates some of the *π*-helices not caught in DSSP and STRIDE assignment programs. DISICL[13], focuses on main chain dihedral angles for structure assignment. SEGNO[14] generates more reliable assignments for distorted locations in the structure.

DEFINE[15], P-CURVE[16], PROSIGN[17], P-SEA[18], PALSSE[19], STICK[20], VoTAP[21] and SABA[22] uses atomic coordinate information for structure assignment wherein SABA introduces the concept of pseudo center, an imaginary point that lies between two consecutive *C*_*α*_ atoms. VoTAP[21] applies Voroni tessellation to establish contacts between residues. SKSP[23] performs structural alignment of protein pairs prior to structure assignment, and the similarities with these aligned residues are used for structural assignments. SACF[24] introduces fragment-based approaches for conquering structure assignment problems. Here, assignments are done by aligning *C*_*α*_ fragments against some template fragments. SST[25] implements a Bayesian method that attempts to maximize the joint probability of a hypotheses(regarding secondary structure assignment of given coordinates) and the data. In addition to those mentioned above there are several other related works[26],[27], [28], [29], [30] in this area.

PCASSO[31], a machine learning-based structure assignment program achieves significant improvements in accuracy and speed compared to other state-of-art methods. PCASSO applies random forest, a supervised learning approach for feature classification. Using *C*_*α*_ and pseudo center coordinates, a feature vector consisting of 258 feature elements were calculated for each of the residues. These features are then processed using decision trees to assign structures to the residues. The best split for each node in the tree uses 16 random features out of 258 features. This method, based solely on *C*_*α*_ coordinates, shows high speed and accuracy when compared to those that require intensive bond calculations and coordinate geometry analysis.

Random forest secondary structure assignment(RaFoSA)[32], another Artificial Intelligence (AI) technique, learns the secondary structure details using a random forest classifier and assigns a residue to one of the secondary structure classes. Features used by RaFoSA include residue type, *C*_*α*_−*C*_*α*_ distances, angle between three *C*_*α*_ atoms, torsional angle formed by four *C*_*α*_ atoms and residue-residue contacts. This method finds its applicability in coarse-grained as well as all atom-based protein systems.

The above studies show that individual relationships among 3D coordinates of a protein structure clearly distinguish various secondary structure elements. Recent years witnessed a considerable deposition of data in protein repositories, and this abundance of data always enhance the performance of Deep Learning(DL) algorithms. DL technologies coupled with big data and GPU accelerated computing has made a broader impact in many areas, including protein modelling and design[33][34][35][36][37]. Utilizing the sheer volumes of data combined with DL techniques, it is now possible to extract the relationships existing in the input data to divert that knowledge to structure assignment problems. Still, the works in structure assignment using DL techniques is very rare or almost none. Here, a Convolutional Neural Network(CNN) based model automates structure assignment process. As protein fragments are easier to handle than full-length proteins, and to assess DL techniques’ applicability to structure assignment process, the paper’s model works with fragment structure assignments. The model takes protein fragments from homosapiens labelled by their three-state secondary structure information from protein structure file(PDB) files. Even though there are many direct and accurate methods for assigning secondary structure from tertiary protein structures, a DL based solution that gives faster results and better accuracy will benefit the computational biologists working on protein learning tasks.

The paper is organized as follows: Section 1 introduces and describes some of the state-of-art methods. Section 2 outlines the DLFSA design. Section 3 explains the dataset construction and other implementation aspects. Section 4 presents the results and discussions. Section 5 concludes the work.

## 2 Methods

Deep Learning-based Fragment Structure Assignment(DLFSA) is a deep learning model that predicts the secondary structures of protein fragments from atomic coordinate representations. The model outputs three types of secondary structures, viz. Helices(H), Sheets(E) and Coils/others(C). Since there is no standard fragment structure assignment dataset available for training these deep learning models, the design process’s first step is constructing a fragment library. A subset of these fragments are prepossessed and fed into the proposed CNN model. CNN[38] are a class of deep neural networks that are suitable for learning the patterns from a sequence of *C*_*α*_ coordinates and predicts whether the sequence forms a helix, sheet or coil. Fig. 1 shows a high-level design of the proposed work.

**Fig. 1:**
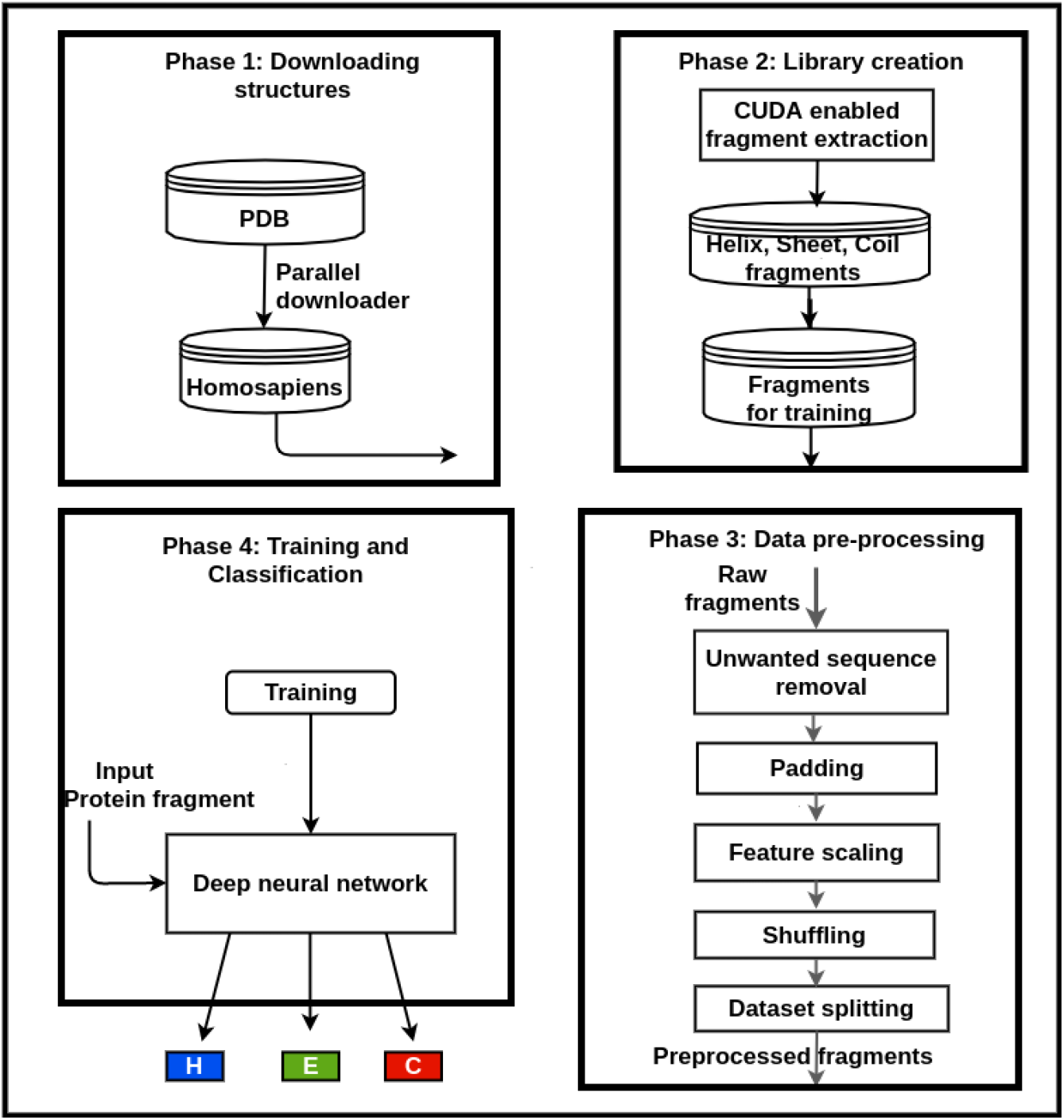
Workflow of DLFSA

### 2.1 CNN for structure assignment

CNN are neural networks that extract relevant features from input data points by applying several filters. CNN[36][39][40][41] are quite powerful at capturing relationships among spatial data. Protein fragments represent a continuous sequence of coordinates in space. Each input file in the training set consists of 9 × 4 array on which the convolution window moves to extract the features. The CNN starts with a kernel window of size 2 × 2 followed by 16 filters and 64 filters. Max pooling technique performs dimensionality reduction followed by four fully connected layers. Dropout regularization method prevents overfitting. The classifier model ends in three nodes with a softmax activation function(equation 2) that predicts a structure’s probability to a helix, sheet or coil. Labels were *one hot* encoded, with 0, 1, 2 representing sheet, helix and coil respectively. Fig. 2 shows the proposed CNN model’s architecture with all its layers, filters, and other parameters.

**Fig. 2:**
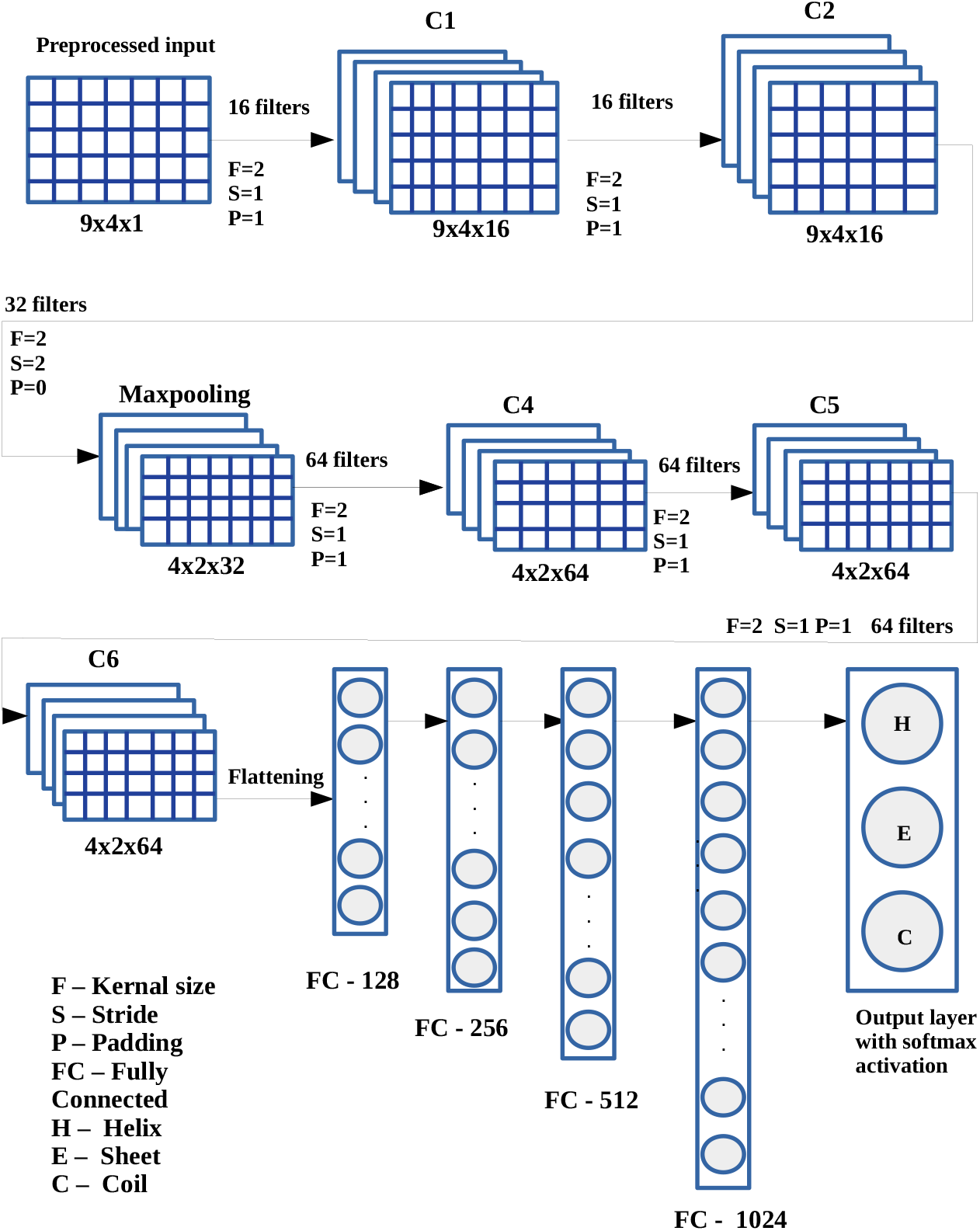
CNN model summary

The softmax function is given by the equation:

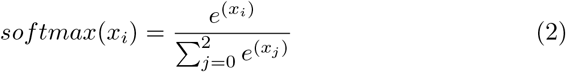

### 2.2 Dataset generation

3D protein structures of 42,000 homosapiens from PDB[42] repository contributes to the construction of the fragment library. A PDB file includes much information regarding the experimental structure of a protein. The *C*_*α*_ coordinates of the downloaded protein files were extracted and stored in a local repository. Each line in a PDB file is called a record. An ATOM record in a PDB file holds the atomic coordinate information of residues involved in protein formation.

The secondary structure information available in the PDB files provides labels to the constructed fragment library. CUDA implements GPU level parallelism to the fragment extraction codes written in C++. This parallel program generates protein fragments within an acceptable time frame. Algorithm 1 explains the pseudocode for extracting the helix and sheet information from PDB files. After executing Algorithm 1, varying length helical, sheet and coil segments, and other essential details, are collected. This secondary structure information is used as input by Algorithm 2 for extracting fixed-sized fragments.

Separate directories are created for different fragments, namely *Helix*, *Sheet* and *Coil* and for different fragment sizes. A kernel function and some host functions offload parallel data operations to the GPU device. Even though the algorithm can extract fragments of any length, the proposed work uses only three, five, six and nine length fragments. Algorithm 2 explains the pseudocode for creating the fragment libraries of desired sizes, and Table 1 shows the fragment library details. Algorithm 3(Supplementary materials) outlines the kernel function.

**Algorithm 1.**
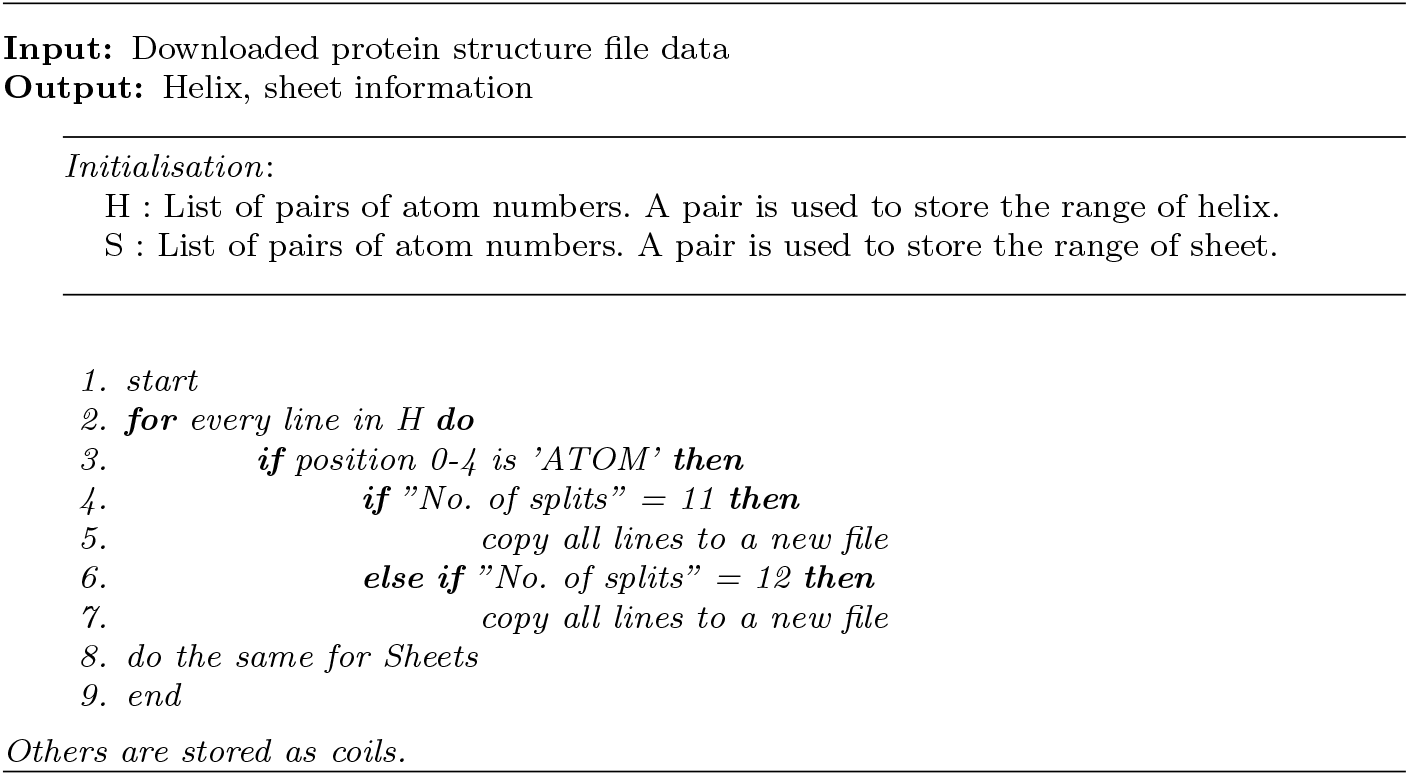
Pseudocode for extracting helix and sheet information

**Table 1:** Fragment library

| Fragment type | Length | No. of fragments | No. of residues |
| --- | --- | --- | --- |
| Helix | 9 | 21 844 | 1 96 596 |
| Sheet | 9 | 16 159 | 1,45 431 |
| Coil | 9 | 4 70 591 | 42 35 319 |
| Helix | 6 | 1 16 938 | 7 01 628 |
| Sheet | 6 | 19 030 | 1 14 180 |
| Coil | 6 | 6 89 466 | 41 36 796 |
| Helix | 3 | 29 26 924 | 87 80 772 |
| Sheet | 3 | 59 572 | 1 78 716 |
| Coil | 3 | 13 23 127 | 39 69 381 |

A fragment is categorized as a helix in the fragment extraction process only if all residues took part in helix formation. The fragments are not just approximations, but exact categorization of its secondary structure. The output(label) indicates the secondary structure information of a fragment as a whole. Approximating a fragment structure class based on its majority residue nature is common in fragment library creation algorithms[43](e.g. if four residues out of six are involved in a helix formation, then it is categorized as a helix).

The proposed classification method considers all fragments other than the helices and sheets as coils. So, coil fragment libraries contain more fragments compared to others. These imbalances in the counts of helices, sheets and coils create problems while training the model. The model training chooses a subset of fragments with a similar count. Table 2 shows the fragment count statistics.

**Algorithm 2.**
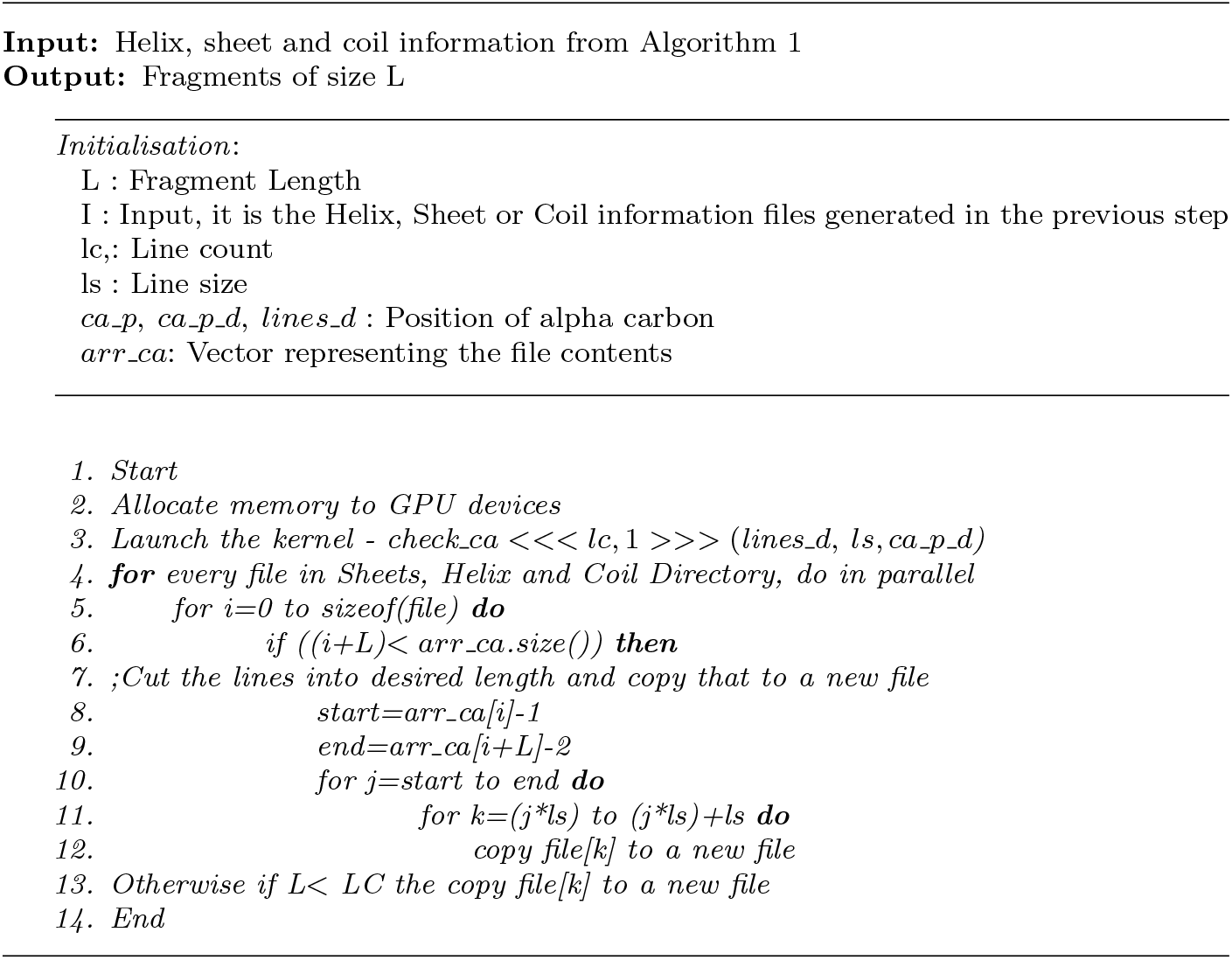
Pseudocode for creating a fragment library of length L

**Table 2:**
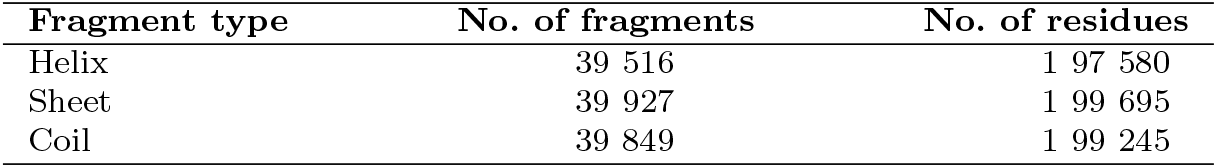
Data sets used

#### 2.2.1 Data preprocessing

The data set consists of 1,19,281 fragment files containing residue names, atom names, atomic coordinates, occupancy factors, Etc. From these files, atomic coordinates of C*α* atoms(7,15,671 C*α* coordinates in total, Table 3), along with corresponding residue names are extracted and stored for further processing. The preprocessing step considers only fragments of length nine, six, five and three. Fragment library construction for fragment assembly methods selects mixed length fragments of size three, six, and nine as a complete set. The fragments under consideration are equalized to the length of available maximum, by padding zeros in the end. Next step converts the fragments into an array of dimension 9*X*4, where nine is the maximum allowable fragment length and, each line in a fragment file contains a residue name along with three coordinates of C*α* atom. The integers from 0 to 19 represent the residue names in the dataset. The final step scales the input features to [−1,1] for better convergence of the model. Finally, the data set is randomly shuffled and split for training, testing and validation purposes. Fig. 1:Phase 3 shows the steps involved in data preprocessing.

**Table 3:**
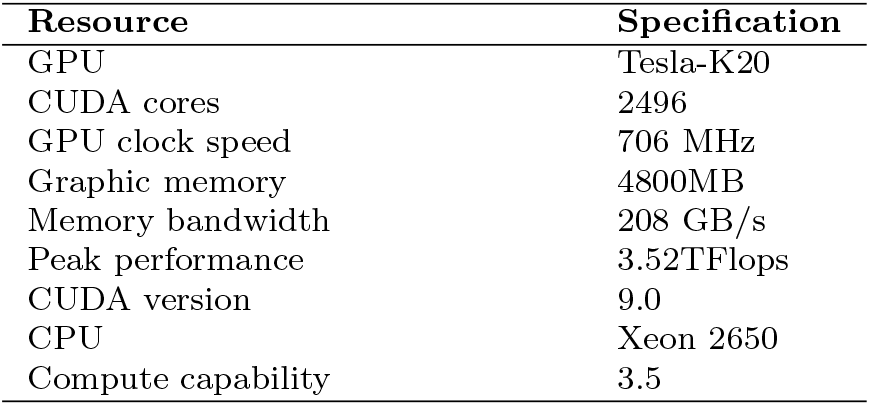
Hardware configuration

| Resource | Specification |
| --- | --- |
| GPU | Tesla-K20 |
| CUDA cores | 2496 |
| GPU clock speed | 706 MHz |
| Graphic memory | 4800MB |
| Memory bandwidth | 208 GB/s |
| Peak performance | 3.52TFlops |
| CUDA version | 9.0 |
| CPU | Xeon 2650 |
| Compute capability | 3.5 |

### 2.3 Hardware configuration

Table 3 shows the hardware configuration of the system used for fragment library creation.

#### 2.3.1 DLFSA web portal

The fragmented structure assignment model, DLFSA is made available to the public through the web portal - www.proteinallinfo.in. The web interface uses python Django[44] framework. DLFSA takes a fragment sequence as the input, and assign a secondary structure to it. The fragment sequence consists of the residue names, atomic coordinates of its C*α* atoms and other parameters as in the PDB file format. The portal displays a sample file format. The maximum allowable length of a testing fragment is nine. The predicted structure represents the fragment’s structure as a whole. The library generation source codes and the model codes are made open(https://github.com/jisnava/DLFSA/).

## 3 Results and Discussion

The results and accuracy of DLFSA, when compared to state-of-art methods, are given below.

### 3.1 Timing analysis

The fragment extraction algorithm written in CUDA C++ executes parallelly on the downloaded PDB files of homosapiens. The algorithm executed on Tesla K20 GPU card achieves significant speed enhancements over its serial counterpart. Fig. 4 shows the comparison statistics. The overall speedup achieved is given in Table 4, where *T*_*s*_ is the serial execution time and *T*_*p*_ is the parallel execution time.

**Fig. 3:**
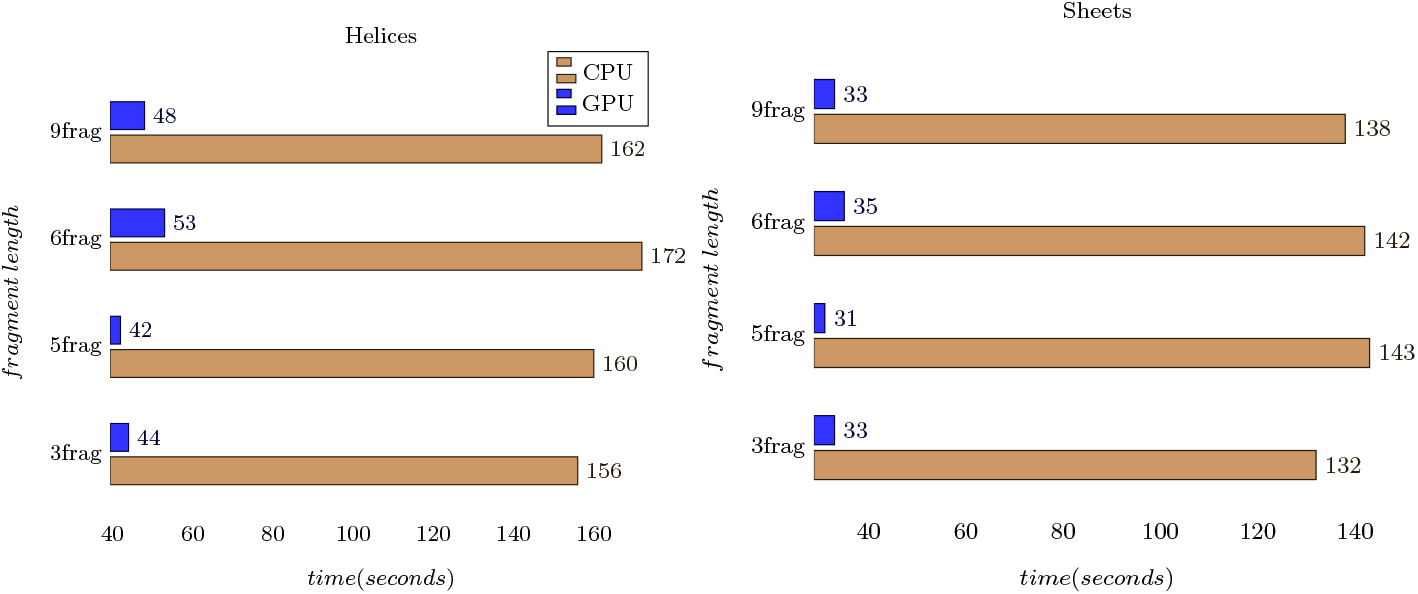
CPU and GPU timing analysis plots for helix and sheet structures

**Fig. 4:**
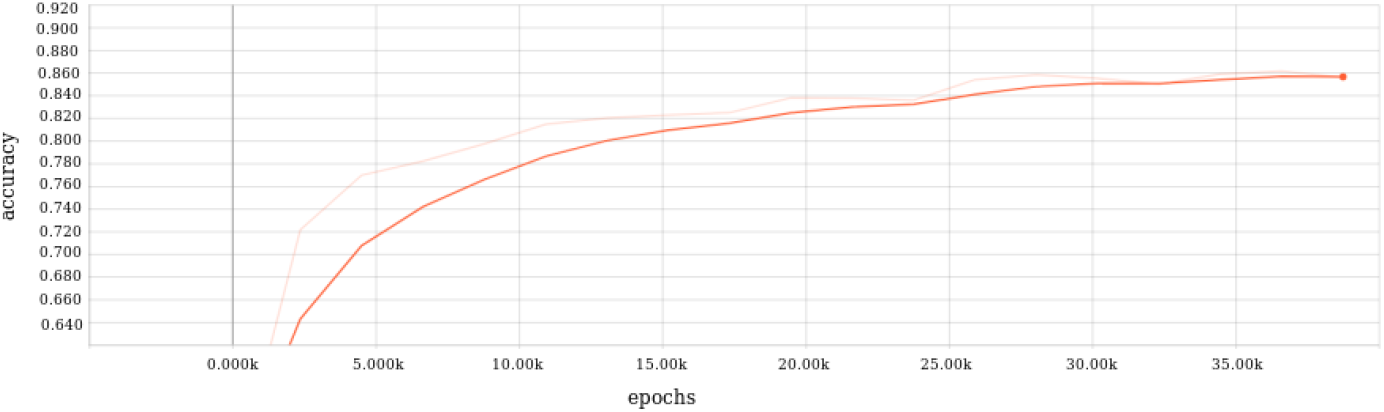
Plot showing the model accuracy

**Table 4:** Speedup achieved

| Fragment type | $T_s$(seconds) | $T_p$(seconds) | Speedup($T_s/T_p$) |
| --- | --- | --- | --- |
| Helix | 11 215 | 39 007 | 28.75 |
| Sheet | 8 081 | 33 351 | 24.23 |

### 3.2 Accuracy

The proposed model uses TensorFlow-Keras framework. Input features to the neural network take a batch size of 512. The activation function used for convolution is ReLU, and softmax for classification. The model is tuned to a learning rate of 0.001 and achieved a train and test accuracy of 88.1% and 82.5% respectively. Fig. 5 shows the accuracy graph and Fig. 6 plots the training and validation losses. The random selection of libraries’ fragments may result in 5.6% difference in train and test accuracy. Since these fragments do not evenly distribute in all coordinate ranges, there is a chance of bias towards a range of coordinate values that results in miss-predictions of some coordinates. By applying techniques like clustering, fragments can be grouped based on some distance measures so that the cluster representatives cover a range of coordinate values.

**Fig. 5:**
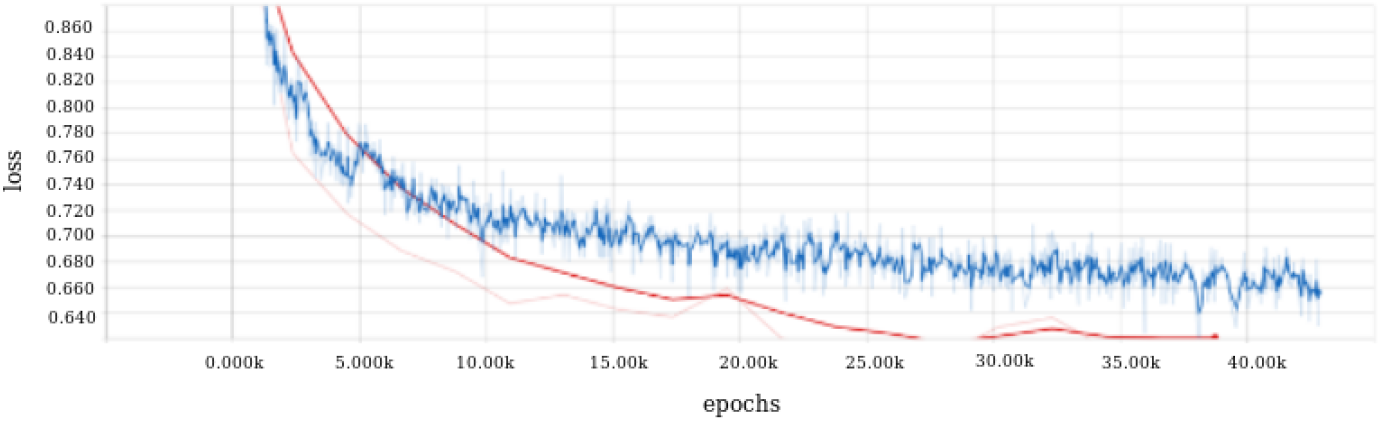
Plot showing training [Blue] and validation losses [Red]

**Fig. 6:**
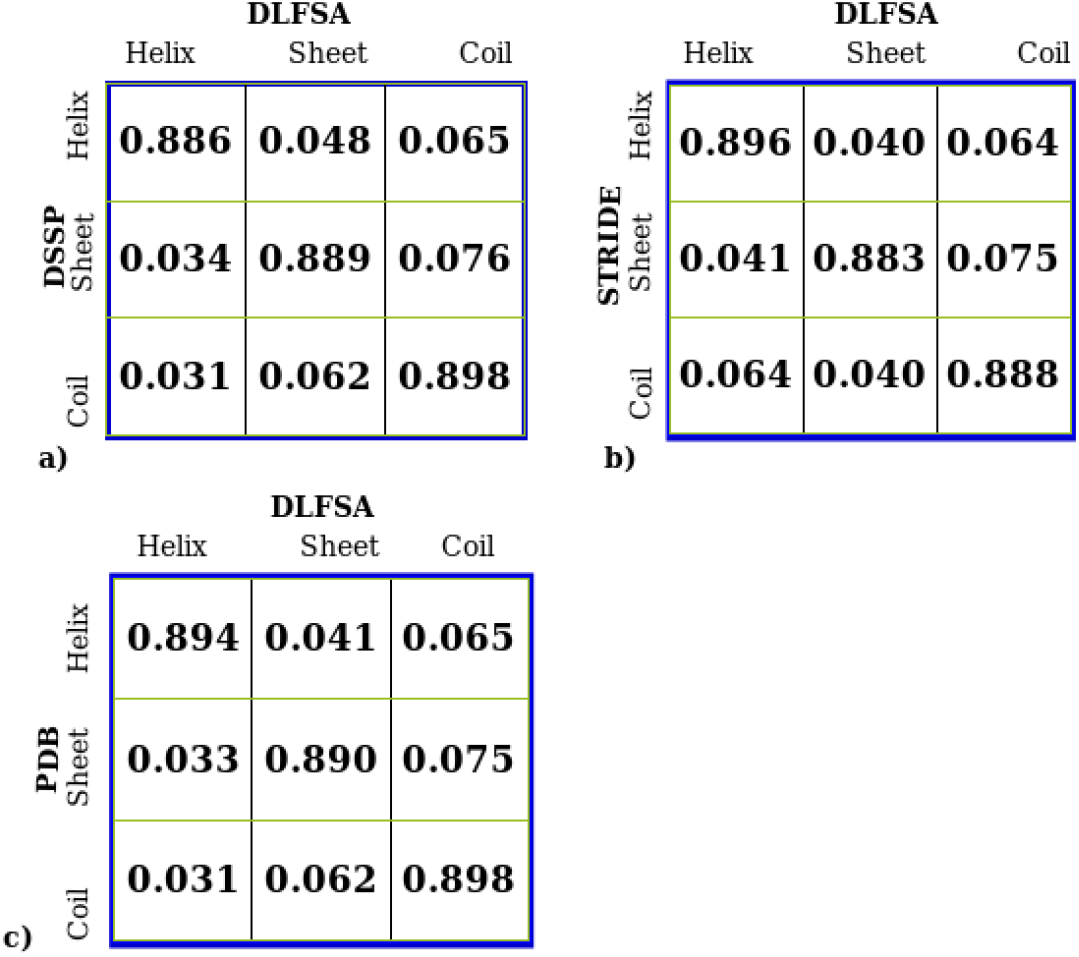
Agreements between DLFSA and other methods. 7(a)(b)(c) Comparison of DLFSA with DSSP, STRIDE and PDB respectively, for a set of randomly chosen fragments

As there are more than 30 methods available in protein structure assignment techniques, it is practically impossible to compare the results with each of them. There are difficulties in accessing the URL mentioned in the papers, and some source codes we could manage to download are not executable, due to unresolved dependencies and lack of documentation. Hence results from DLFSA is compared with the tools which are considered as the gold standards, viz. DSSP, STRIDE and also with the PDB data and some recent methods. The commonly used mapping M:(HGIEBTS-) → (hhhssccc) applies eight-state to three-state secondary structure reductions. The comparison chooses three-set of fragments for which DSSP assigns helix, Sheet and coil structures respectively. Fig. 7 tabulates the results of DLFSA program execution against DSSP, STRIDE and PDB. The details of fragments(including fragment length, protein identifier, chain name and starting position) used to compare with DSSP are provided in additional materials. Comparison with STRIDE and PDB also chooses a similar set of fragments.

**Fig. 7:**
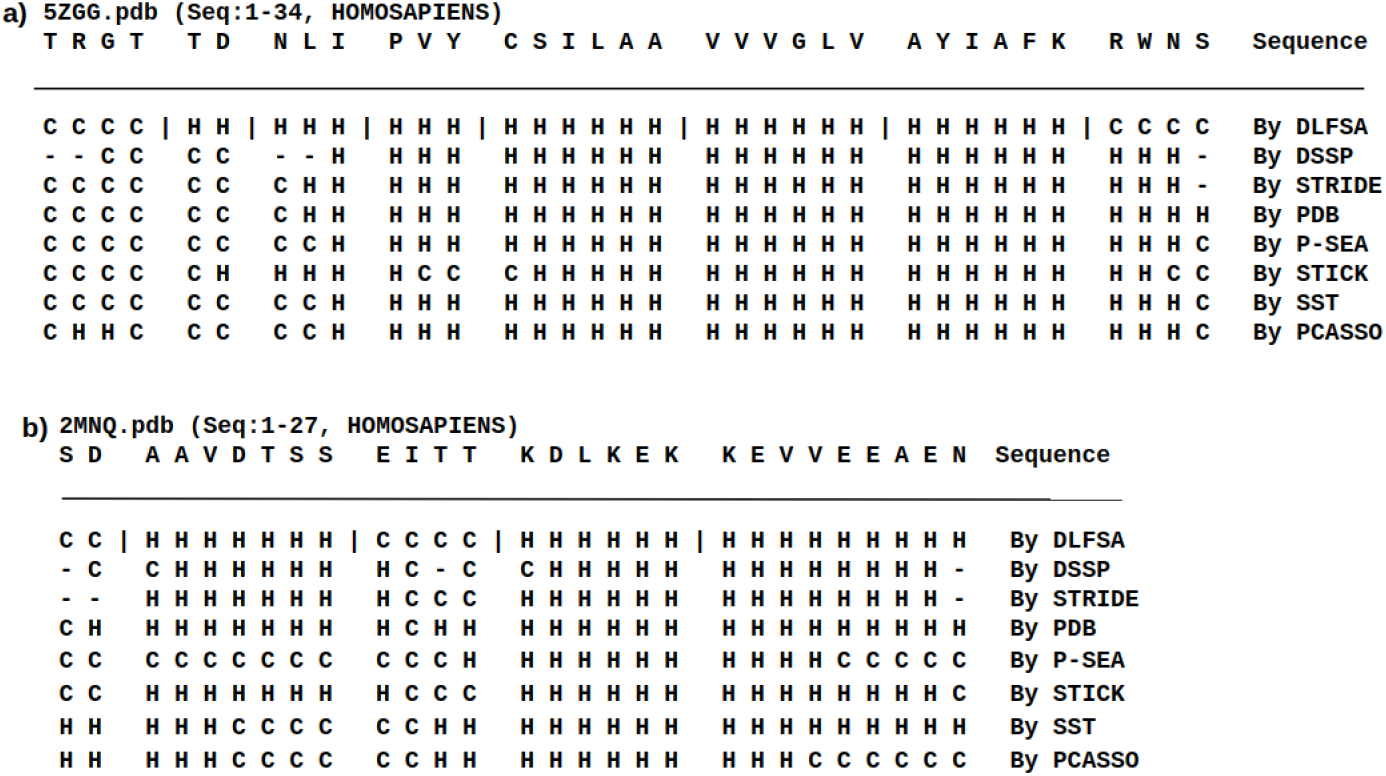
Secondary structure assignments by DLFSA, DSSP, STRIDE, PDB, P-SEA, STICK, SST and PCASSO for two proteins 5zgg(8a) and 2nmq(8b). The first row of the figure shows the primary sequence. The remaining rows represents the secondary structure assignments done by various methods. The symbol separates the non-overlapping random fragments taken for comparison.

Even though the experiment uses limited length protein fragments, its applicability extends to small-length proteins. For full-length protein structures, the analysis uses a combination of varying length non-overlapping random fragments. The method successfully predicts the secondary structure information for small proteins. Fig. 8 shows the results of applying DLFSA on proteins from homosapiens, and Fig. 9 shows the secondary structure assignments done on a protein(ID: 2mnq), by various programs visualized through Chimera.

**Fig. 8:**
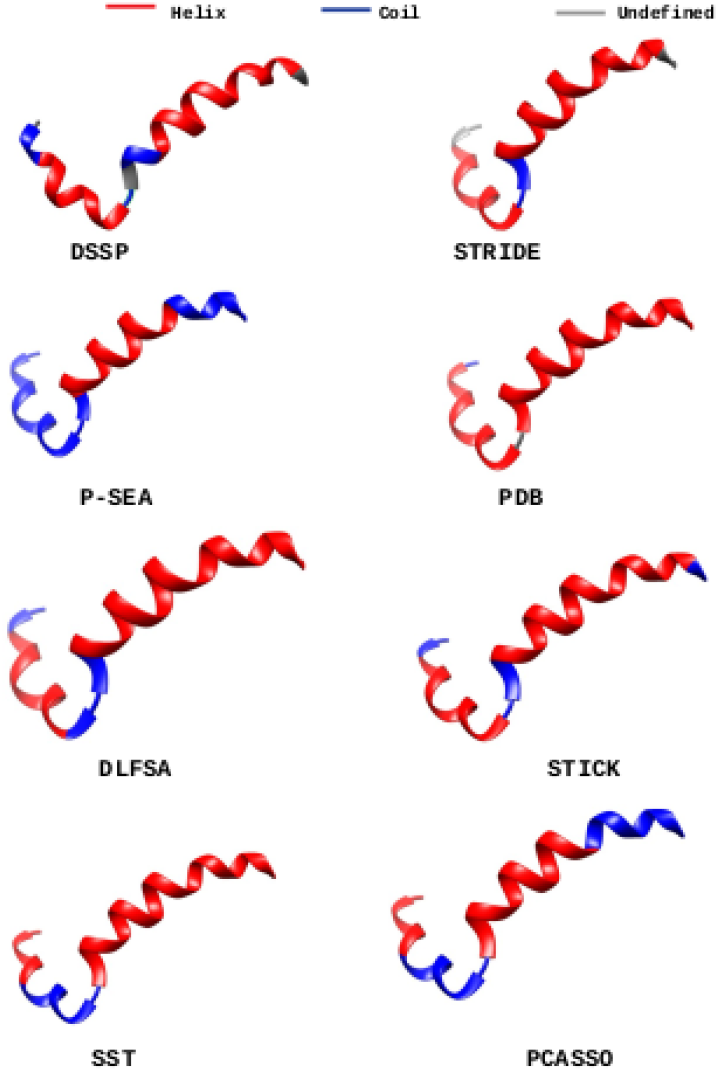
Secondary structure assignments by DLFSA, DSSP, STRIDE, P-SEA, PDB, SST and PCASSO for protein 2nmq.

The results obtained shows that the proposed model gives better results on secondary structure assignments of protein fragments, using the features extracted from Cartesian coordinates. The model has been implemented for three state secondary structures, viz. helices, sheets and coils to enable smooth learning. The main difficulty in implementing the model for eight-state secondary structures is that all the eight states may not occur with equal probability in protein structures and hence, the model will not learn consistently from such unbalanced data. The same problem also occurs in secondary structure prediction tasks, when trained with the primary protein sequences and its eight-state secondary structure information. The fragment-based model implemented in the paper can be benefited to homology modelling systems also. These DL models for structure assignments(when extended to full-length, multi-domain proteins) finds application in quality assessment stages of protein structure prediction systems.

## 4 Conclusion

With the increase of 3D structures in protein repositories, it is now possible to automatically assign protein structures without human intervention. Today, state-of-the-art systems use machine learning with manually engineered feature extraction, and none of the assignment systems currently available is entirely based on Deep Learning techniques. We developed a CNN based model to automate protein structure assignment process. The model learns the spatial relationships among protein coordinates and utilizes it for secondary structure assignments. The model is successfully tested on protein fragments and a few full-length proteins. Assignment systems able to extract local and global features from the protein structures and use this to guide the structure assignment process itself, are now possible with Deep Learning. When provided with sufficient data, deep learning models outperform traditional approaches, in Natural Language Processing, Computer Vision, and Speech Recognition Systems, to name a few. Our model accuracy unarguably validates its applicability to secondary structure assignment problems and in challenging environments where only *C*_*α*_ atoms are available. The developed model shows comparable accuracy with the two gold standard methods in the area. This experiment highlights neural networks’ ability to capture local structures from coordinate data. The current model extends to include 8-state secondary structures(Q8) for more precise predictions, provided with enough experimental data. Further studies are needed to develop this technique to large and multi-domain protein structures. With increasing computational powers and experimental data, more improvements are expected from computational biologists, on protein structure assignment tasks, for a faster and more accurate solution.

## Declarations

### Availability of data and material

DLFSA is made available to the public through the web portal - www.proteinallinfo.in. The datasets generated during and/or analysed during the current study are not publicly available due to its large size but are available from the corresponding author on reasonable request.

### Code availability

The library generation source codes and the model codes are made open at https://github.com/jisnava/DLFSA/.

### Authors Contributions

V. A. Jisna(VAJ) did the conceptualization. Prayagh Madhu did the model coding. VAJ developed the web portal and wrote the manuscript. P. B. Jayaraj supervised the project. All authors read and approved the final manuscript.

## Acknowledgments

Authors would like to thank the Central Computing Centre, National Institute of Technology Calicut(NITC) for providing GPU servers for this work. The authors would like to acknowledge the valuable suggestions from Hemant Yadav in constructing the fragment library and Jinto Antony for developing the web portal.

## Supplementary Materials

**Fig. 9:**
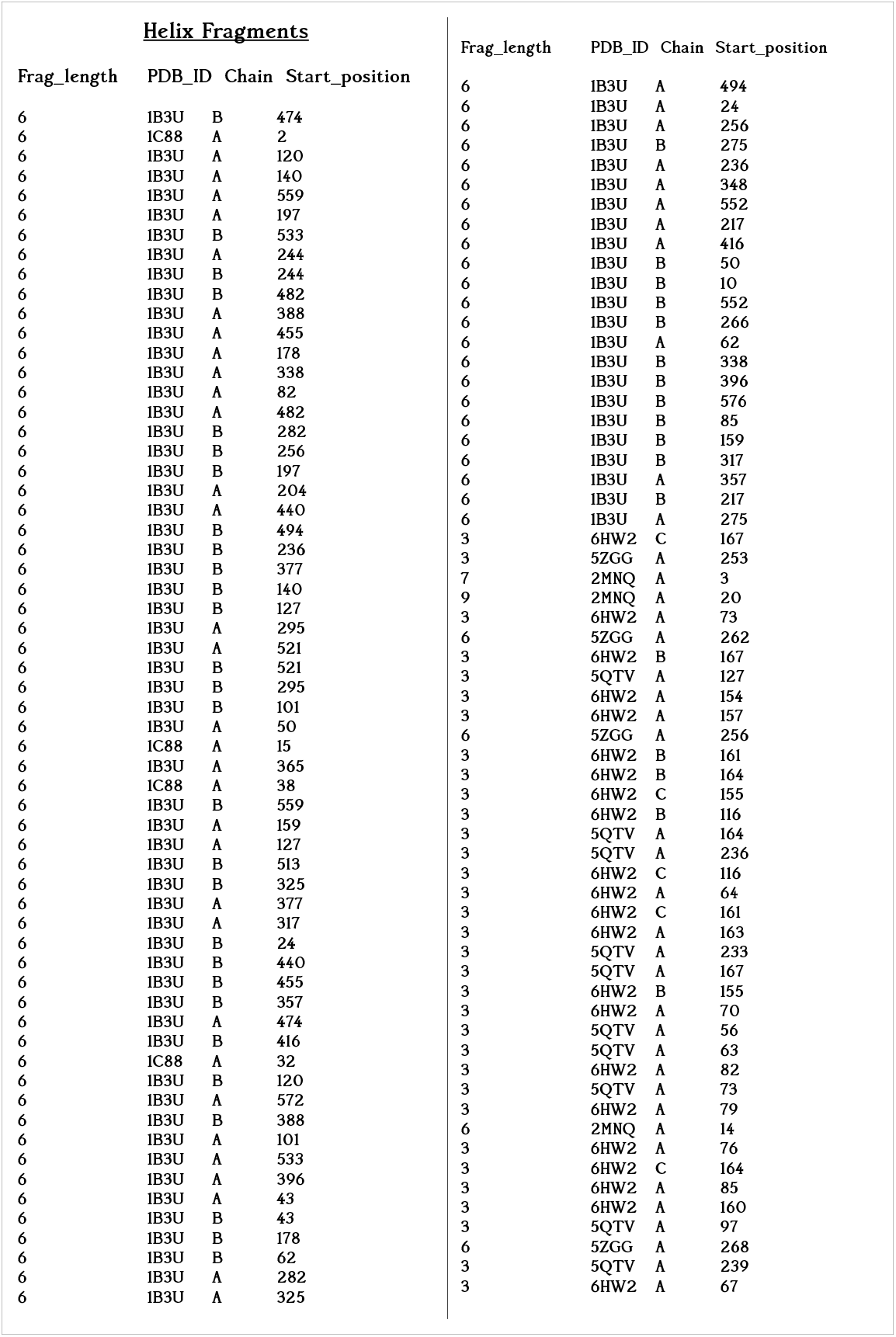
Fragment lengths, PDBID, chain names and fragment starting position of helix fragments used for DLFSA - DSSP comparison.

**Fig. 10:**
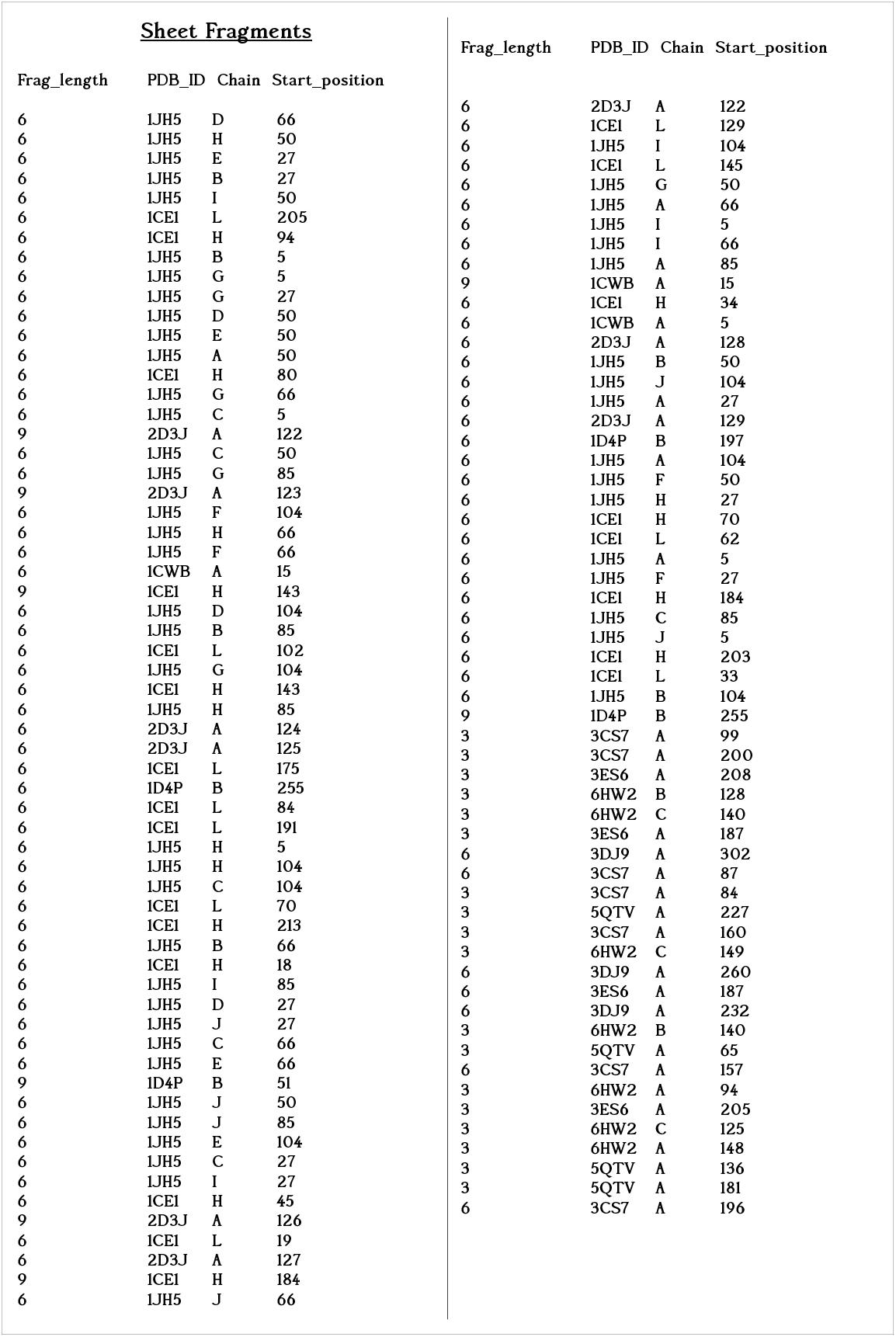
Fragment lengths, PDBID, chain names and fragment starting position of sheet fragments used for DLFSA - DSSP comparison.

**Algorithm 3.**
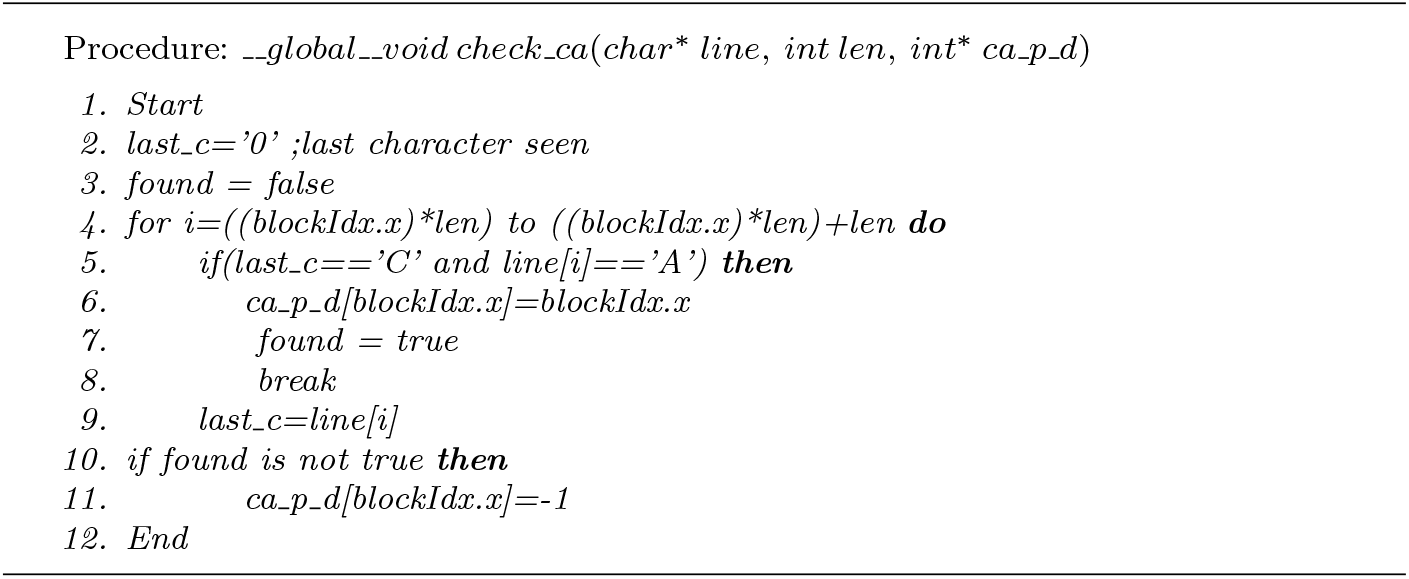
Kernel function for finding c*α* atom position

## References

1. L. Pauling, R. B. Corey, H. R. Branson, The structure of proteins: two hydrogen-bonded helical configurations of the polypeptide chain, Proceedings of the National Academy of Sciences 37 (4) (1951) 205–211.

2. J. Reeb, B. Rost, Secondary structure prediction, Encyclopedia of Bioinformatics and Computational Biology (2019) 488–496.

3. R. Srinivasan, G. D. Rose, A physical basis for protein secondary structure, Proceedings of the National Academy of Sciences 96 (25) (1999) 14258–14263.

4. D. Eisenberg, The discovery of the *α*-helix and *β*-sheet, the principal structural features of proteins, Proceedings of the National Academy of Sciences 100 (20) (2003) 11207–11210.

5. J. Zhou, H. Wang, Z. Zhao, R. Xu, Q. Lu, Cnnh pss: protein 8-class secondary structure prediction by convolutional neural network with highway, BMC bioinformatics 19 (4) (2018) 60.

6. J. Abbass, J.-C. Nebel, N. Mansour, Ab initio protein structure prediction: methods and challenges, Biol Knowl Discov Handb. Hoboken, New Jersey: John Wiley & Sons, Inc (2013) 703–24.

7. C. B. Anfinsen, Principles that govern the folding of protein chains, Science 181 (4096) (1973) 223–230.

8. J. N. Onuchic, P. G. Wolynes, Theory of protein folding, Current opinion in structural biology 14 (1) (2004) 70–75.

9. W. Kabsch, C. Sander, Dictionary of protein secondary structure: pattern recognition of hydrogen-bonded and geometrical features, Biopolymers: Original Research on Biomolecules 22 (12) (1983) 2577–2637.

10. G. t. Ramachandran, V. Sasisekharan, Conformation of polypeptides and proteins, in: Advances in protein chemistry, Vol. 23, Elsevier, 1968, pp. 283–437.

11. J. Zacharias, E.-W. Knapp, Protein secondary structure classification revisited: processing dssp information with pssc, Journal of chemical information and modeling 54 (7) (2014) 2166–2179.

12. M. Fodje, S. Al-Karadaghi, Occurrence, conformational features and amino acid propensities for the *π*-helix, Protein Engineering, Design and Selection 15 (5) (2002) 353–358.

13. G. Nagy, C. Oostenbrink, Dihedral-based segment identification and classification of biopolymers i: proteins, Journal of chemical information and modeling 54 (1) (2014) 266–277.

14. M. V. Cubellis, F. Cailliez, S. C. Lovell, Secondary structure assignment that accurately reflects physical and evolutionary characteristics, BMC bioinformatics 6 (S4) (2005) S8.

15. F. M. Richards, C. E. Kundrot, Identification of structural motifs from protein co-ordinate data: secondary structure and first-level supersecondary structure, Proteins: Structure, Function, and Bioinformatics 3 (2) (1988) 71–84.

16. H. Sklenar, C. Etchebest, R. Lavery, Describing protein structure: a general algorithm yielding complete helicoidal parameters and a unique overall axis, Proteins: Structure, Function, and Bioinformatics 6 (1) (1989) 46–60.

17. S.-R. Hosseini, M. Sadeghi, H. Pezeshk, C. Eslahchi, M. Habibi, Prosign: A method for protein secondary structure assignment based on three-dimensional coordinates of consecutive c*α* atoms, Computational biology and chemistry 32 (6) (2008) 406–411.

18. G. Labesse, N. Colloc’h, J. Pothier, J.-P. Mornon, P-sea: a new efficient assignment of secondary structure from c*α* trace of proteins, Bioinformatics 13 (3) (1997) 291–295.

19. I. Majumdar, S. S. Krishna, N. V. Grishin, Palsse: A program to delineate linear secondary structural elements from protein structures, BMC bioinformatics 6 (1) (2005) 202.

20. W. R. Taylor, Defining linear segments in protein structure, Journal of molecular biology 310 (5) (2001) 1135–1150.

21. F. Dupuis, J.-F. Sadoc, J.-P. Mornon, Protein secondary structure assignment through voronoi tessellation, Proteins: structure, function, and bioinformatics 55 (3) (2004) 519–528.

22. S.-Y. Park, M.-J. Yoo, J.-M. Shin, K.-H. Cho, Saba (secondary structure assignment program based on only alpha carbons): a novel pseudo center geometrical criterion for accurate assignment of protein secondary structures, BMB reports 44 (2) (2011) 118–122.

23. W. Zhang, A. K. Dunker, Y. Zhou, Assessing secondary structure assignment of protein structures by using pairwise sequence-alignment benchmarks, Proteins: Structure, Function, and Bioinformatics 71 (1) (2008) 61–67.

24. C. Cao, G. Wang, A. Liu, S. Xu, L. Wang, S. Zou, A new secondary structure assignment algorithm using c*α* backbone fragments, International journal of molecular sciences 17 (3) (2016) 333.

25. A. S. Konagurthu, A. M. Lesk, L. Allison, Minimum message length inference of secondary structure from protein coordinate data, Bioinformatics 28 (12) (2012) i97–i105.

26. D. Klose, B. A. Wallace, R. W. Janes, 2struc: the secondary structure server, Bioinformatics 26 (20) (2010) 2624–2625.

27. P. Kumar, M. Bansal, Identification of local variations within secondary structures of proteins, Acta Crystallographica Section D: Biological Crystallography 71 (5) (2015) 1077–1086.

28. M. Habibia, C. Eslahchia, H. Pezeshkc, M. Sadeghid, An information theoretic approach to secondary structure assignment (2008).

29. T. Taylor, M. Rivera, G. Wilson, I. I. Vaisman, New method for protein secondary structure assignment based on a simple topological descriptor, Proteins: Structure, Function, and Bioinformatics 60 (3) (2005) 513–524.

30. Y. Zhang, C. Sagui, Secondary structure assignment for conformationally irregular peptides: Comparison between dssp, stride and kaksi, Journal of Molecular Graphics and Modelling 55 (2015) 72–84. doi:https://doi.org/10.1016/j.jmgm.2014.10.005. URL http://www.sciencedirect.com/science/article/pii/S1093326314001648

31. S. M. Law, A. T. Frank, C. L. Brooks III, Pcasso: A fast and efficient c*α*-based method for accurately assigning protein secondary structure elements, Journal of computational chemistry 35 (24) (2014) 1757–1761.

32. E. O. Salawu, Rafosa: Random forests secondary structure assignment for coarse-grained and all-atom protein systems, Cogent Biology 2 (1) (2016) 1214061.

33. J. Wang, H. Cao, J. Z. Zhang, Y. Qi, Computational protein design with deep learning neural networks, Scientific reports 8 (1) (2018) 6349.

34. J. Cheng, A. N. Tegge, P. Baldi, Machine learning methods for protein structure prediction, IEEE reviews in biomedical engineering 1 (2008) 41–49.

35. B. Zhang, J. Li, Q. Lü, Prediction of 8-state protein secondary structures by a novel deep learning architecture, BMC bioinformatics 19 (1) (2018) 293.

36. Y. LeCun, Y. Bengio, G. Hinton, Deep learning, nature 521 (7553) (2015) 436.

37. G. B. Goh, N. O. Hodas, A. Vishnu, Deep learning for computational chemistry, Journal of computational chemistry 38 (16) (2017) 1291–1307.

38. K. O’Shea, R. Nash, An introduction to convolutional neural networks, arXiv preprint arXiv:1511.08458 (2015).

39. A. Busia, J. Collins, N. Jaitly, Protein secondary structure prediction using deep multi-scale convolutional neural networks and next-step conditioning, arXiv preprint arXiv:1611.01503 (2016).

40. R. Zamora-Resendiz, S. Crivelli, Structural learning of proteins using graph convolutional neural networks, bioRxiv (2019) 610444.

41. M. Niepert, M. Ahmed, K. Kutzkov, Learning convolutional neural networks for graphs, in: International conference on machine learning, 2016, pp. 2014–2023.

42. https://www.rcsb.org/structure/1AU1/, accessed : 2020-09-09.

43. J. B. Holmes, J. Tsai, Some fundamental aspects of building protein structures from fragment libraries, Protein science 13 (6) (2004) 1636–1650.

44. https://www.djangoproject.com/, accessed : 2020-12-12.

